# A more expressive spline representation for SBML models improves code generation performance in AMICI

**DOI:** 10.1101/2023.06.29.547120

**Authors:** Lorenzo Contento, Paul Stapor, Daniel Weindl, Jan Hasenauer

## Abstract

Spline interpolants are commonly used for discretizing and estimating functions in mathematical models. While splines can be encoded in the Systems Biology Markup Language (SBML) using piecewise functions, the resulting formulas are very complex and difficult to derive by hand. Tools to create such formulas exist but only deal with numeric data and thus cannot be used for function estimation. Similarly, simulation tools suffer from several limitations when handling splines. For example, in the AMICI library splines with large numbers of nodes lead to long model import times.

We have developed a set of SBML annotations to mark assignment rules as spline formulas. These compact representations are human-readable and easy to edit, in contrast to the piecewise representation. Different boundary conditions and extrapolation methods can also be specified. By extending AMICI to create and recognize these annotations, model import can be sped up significantly. This allows practitioners to increase the expressivity of their models.

While the performance improvement is limited to AMICI, our tools for creating spline formulas can be used for other tools as well and our syntax for compact spline representation may be a starting point for an SBML-native way to represent spline interpolants.

## 1 Background

Mathematical modelling has always been an essential component of the scientific method and, thanks to an increase in available computational power, the complexity of the models has been growing rapidly in the last years [4]. Models are defined in terms of a mathematical function that maps a set of parameters, some of them known and some of them unknown, to an observation (possibly stochastic in nature). For example, such mapping could be defined by a system of ordinary differential equations representing a reaction network in systems biology.

The parameters of a mathematical model are usually real numbers, but they can also be infinite-dimensional objects such as functions [7]. A common use case is that of an input function whose governing law is not known and thus cannot be modelled directly, but which is otherwise necessary to predict the evolution of the other components of the system. When dealing with infinite-dimensional objects computationally, it is necessary to employ discretization methods that replace the original set of possible values with a finite-dimensional one. The higher the dimension of this new set, the more similar the discretized problem will be to the original one.

A popular class of discretization methods for univariate functions consists of interpolation methods. A finite set of points, called nodes, is chosen in the function’s domain and the function is parameterized by its values (and possibly some of its derivatives) at the nodes. The value at a point between nodes is obtained by combining the known values at nearby nodes, usually in a way that preserves the continuity of the function (and of some of its derivatives). Among interpolation methods, a very popular and effective approach is spline interpolation [15]: the function is approximated by a piecewise polynomial which satisfies particular conditions in order to ensure continuity at the nodes. In the simplest case this reduces to piecewise linear interpolation. Next to that, the most popular option is to use cubic polynomials and enforcing continuity up to the first or second derivative, resulting in a cubic spline. Continuity up to the first derivative allows the space of cubic splines to span many natural-looking and plausible choices of input functions for most models. For example, splines are commonly used to model and estimate metabolic fluxes [20,13,16].

Systems biology models are often encoded using the Systems Biology Markup Language (SBML) [10], which allows for a tool-agnostic declarative formulation of the system. The SBML format does not have native support for spline interpolation, and neither do other commonly used alternatives such as CellML [1] and BNGL [8]. However, since arbitrary piecewise functions can be defined in MathML, encoding a spline function is possible by calculating its constituting polynomial segments. Software tools that support the SBML specification can then read and simulate such models correctly. In practice, existing software suffers from several limitations when dealing with complex piecewise functions. For example, COPASI [9] succeeds in computing the sensitivities but is not able to import the SBML file when the spline has more than roughly 40 nodes. On the other hand, libRoadRunner [21,18] simulates the model correctly but does not support adjoint sensitivity analysis, making it less suitable for estimating large numbers of parameters. Finally, the AMICI library for simulation and sensitivity analysis [5] supports adjoint sensitivities, but in the presence of deeply nested piecewise functions it currently requires an extremely long time to generate the C**++** code for the model.

In this paper, we will present an extension to AMICI that allows it to efficiently deal with SBML models containing splines. In Section 2 we briefly present cubic Hermite splines, the spline family we support. Then, we present the workflow for creating and simulating SBML models with splines (summarized in Figure 1). We encode the spline in an SBML model using an assignment rule which, in addition to the MathML piecewise formula for the spline, contains an annotation describing the interpolator in a compact form (Section 3). Such annotated assignment rules can be created using the AMICI Python library from a high-level description of the spline, without requiring the user to derive the piecewise formula for it (Section 4). Finally, models can be imported and simulated using any SBML-compliant tool (Section 5). AMICI parses the annotation and uses it to quickly generate the required C**++** code, while other tools will ignore it and fall back to the MathML expression. We conclude the paper with a real-world application of this workflow (Section 6).

**Fig. 1.**
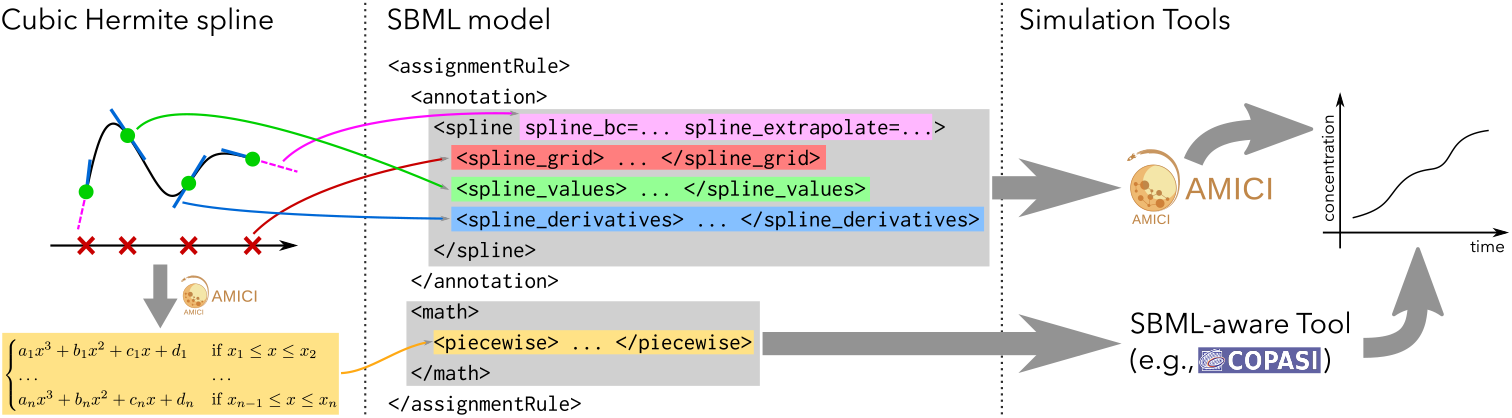
Workflow for adding splines to an SBML model. (i) The user provides a description of a cubic Hermite spline using the nodes positions, the values and derivatives of the spline at the nodes, the boundary conditions and the extrapolation method. (ii) Using the AMICI Python library, an SBML assignment rule for the spline is generated and added to a pre-existing SBML model (the piecewise MathML expression is automatically generated). (iii) The resulting model can be imported and simulated either by AMICI (which parses the annotation) or by other SBML-aware tools (which use the MathML formula).

## 2 Cubic Hermite splines

The space of piecewise cubic polynomials with *n* nodes has 4(*n* − 1) dimensions (each spline has *n* − 1 segments, each completely defined by four coefficients). Different spline families can be obtained by imposing conditions on smoothness and by choosing a standard way to parameterize the remaining degrees of freedom.

In this paper we will be concerned with cubic Hermite splines [15]. Such splines are piecewise cubic polynomials which are continuous up to the first derivative, thus resulting in 2*n* degrees of freedom (two continuity constraints at each inner node, for a total of 2(*n* − 2) constraints). These 2*n* degrees of 4 L. Contento, and P. Stapor, D. Weindl, J. Hasenauer freedom are parameterized by the values of the spline and of its first derivative at all *n* nodes. Often derivatives are computed from the values at the nodes by finite differencing, taking into account the choice of boundary conditions (e.g., enforcing the first or second derivative to be zero), resulting in *n* degrees of freedom. Boundary conditions are deeply linked with the choice of the extrapolation method to use before the first node and/or after the last node. For example, constant extrapolation requires a zero derivative boundary condition to preserve continuity of the first derivative at the boundary. Periodic boundary conditions are also possible, resulting in a dimensionality of 2(*n* − 1) in the case derivatives are specified explicitly, or *n* − 1 when finite differences are used instead. Finally, if the values at the nodes are monotonic, it is possible to ensure monotonicity of the entire spline using appropriate choices for the derivatives at the nodes [3].

However, the most commonly encountered family of cubic splines is that of interpolating cubic splines [15]. These are continuous up to the second derivative and thus have *n* + 2 degrees of freedom, which are parameterized by the spline values at the *n* nodes, plus two boundary conditions. While they are smoother than Hermite splines, the formulas for computing the coefficients for each polynomial piece are more complex. Moreover, the value of the interpolant at any point depends in theory on the values at all nodes, while for a Hermite spline it only depends on the values at the two nearest nodes (excluding the dependence through finite difference approximations of the derivative). This means that, when computing the sensitivities of the interpolated values with respect to the parameters (on which the values defining the spline may depend), using Hermite splines will result in a Jacobian with a more sparse pattern. Finally, it is not possible to enforce monotonicity of the resulting spline, even if the values at the nodes are monotonic.

## 3 Annotation-based compact representation for splines

The native way to represent a cubic Hermite spline in an SBML model is to use a MathML piecewise construct containing the equations of all polynomial segments of the spline. The resulting formula is extremely long and difficult to understand and/or modify. Thus, we propose a higher-level compact XML syntax to specify such splines using their natural parameters: node positions, spline values at the nodes, and either derivatives at the nodes or boundary conditions in case finite differences are to be used. A simple example of one such compact XML representation can be found in Figure 2. Details about the syntax are provided in the Appendix.

**Fig. 2.**
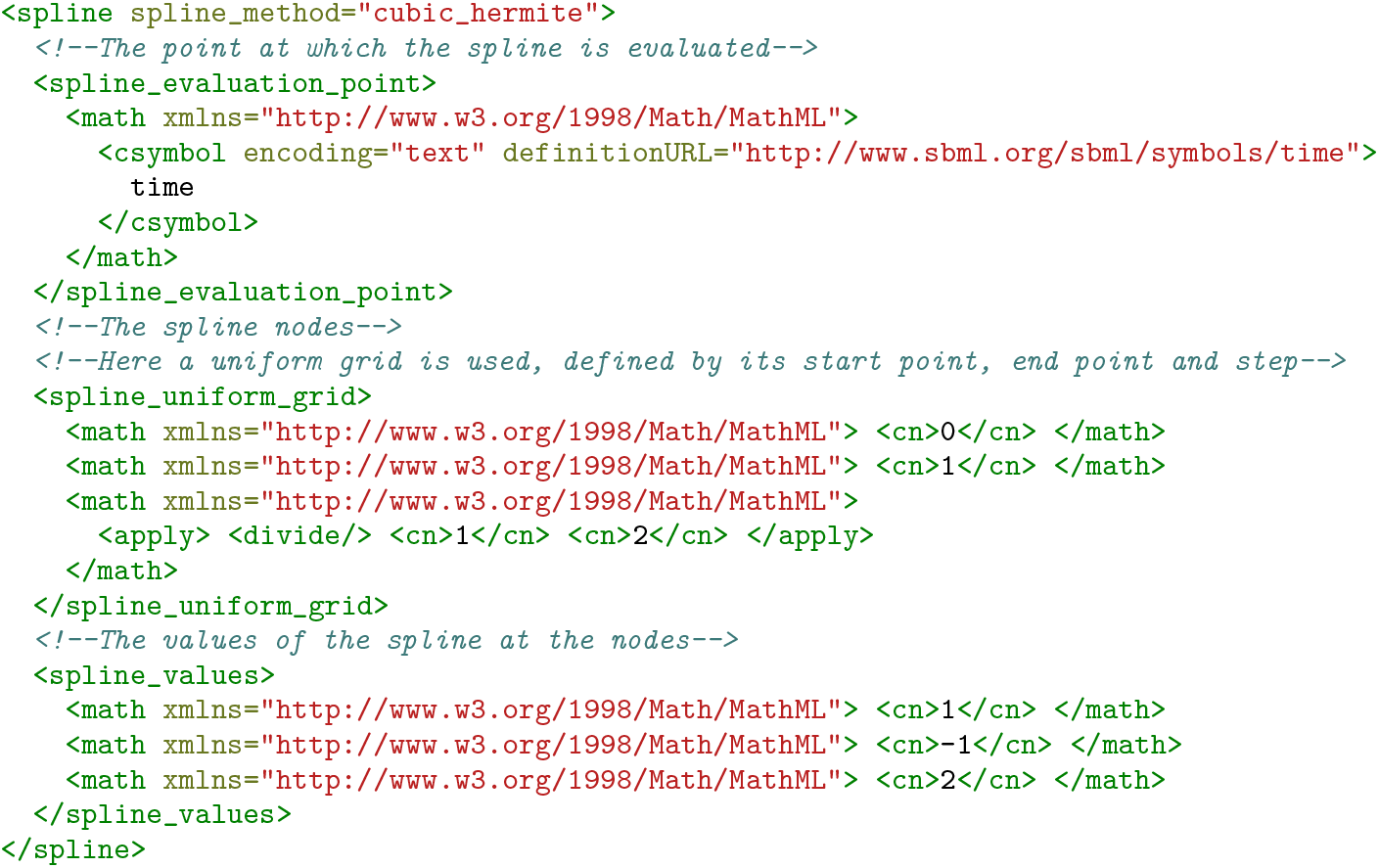
Example of compact XML representation of a cubic Hermite spline with nodes (0, 1*/*2, 1), values at the nodes equal to (1, −1, 2), derivatives computed by finite differences (central formulas for the inner nodes, one-sided formulas for the boundary nodes), and evaluated at the current model time.

## 4 User-friendly insertion of splines into SBML models

A MathML spline representation can be inserted into any other MathML formula appearing in an SBML model. However, for readibility’s sake, one may wish to create a new non-constant parameter for the spline value and create an assignment rule for it, thus avoiding duplication of the very long MathML piecewise formula. This second approach also offers a natural place to store our compact spline representation: the annotation element of the assignment formula. The basic structure of the resulting assignment rule is shown in Figure 3.

**Fig. 3.**
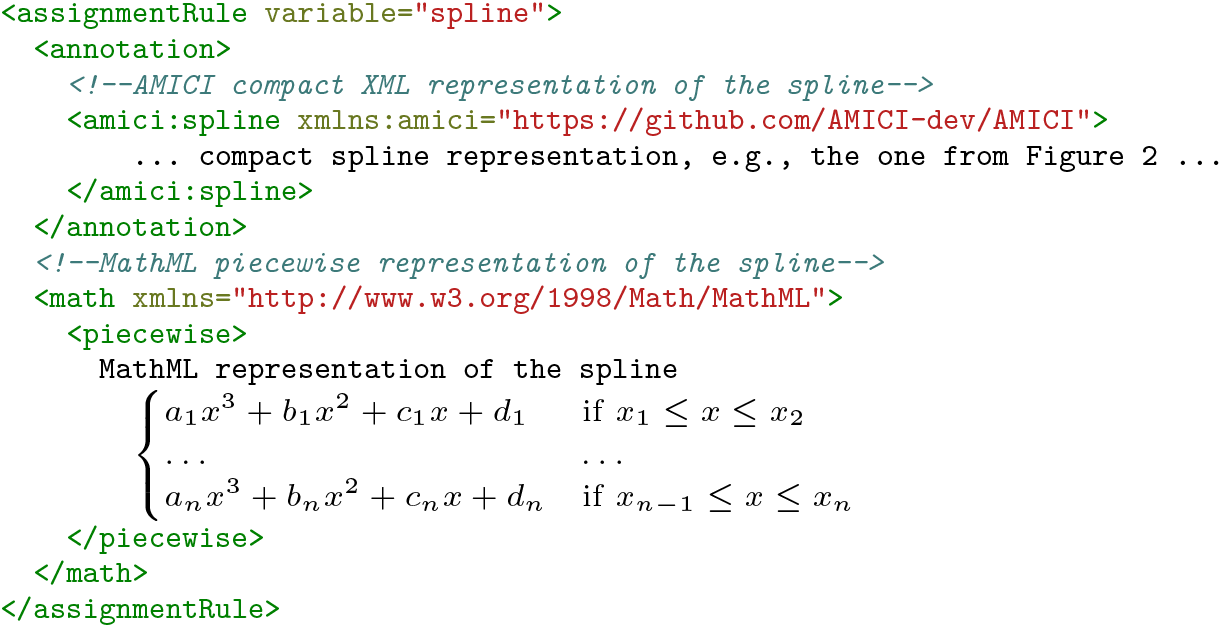
The skeleton of an annotated assignment rule for a spline, including both the AMICI compact representation and the usual MathML piecewise expression.

Since MathML representations of splines are too cumbersome to write manually, they are usually generated with dedicated software tools, such as sbmlutils [11] and SBMLDataTools [14]. However, such tools require the data to be interpolated to be numeric. While very useful for interpolating data series [12], they cannot be used for parameter inference on functions. We have thus added to the AMICI Python library some convenience functions to generate the above mentioned assignment rules, containing both the AMICI-specific compact representation and the usual MathML formula, also covering the case where the spline values at the nodes (or its derivatives) depend on the model parameters. As an example, the code required to add the spline of Figure 2 to an SBML model is shown in Figure 4. More complex spline definitions, including derivatives, boundary conditions and/or extrapolation methods, can be specified; we point the interested reader to the AMICI documentation for the details and various examples.

**Fig. 4.**
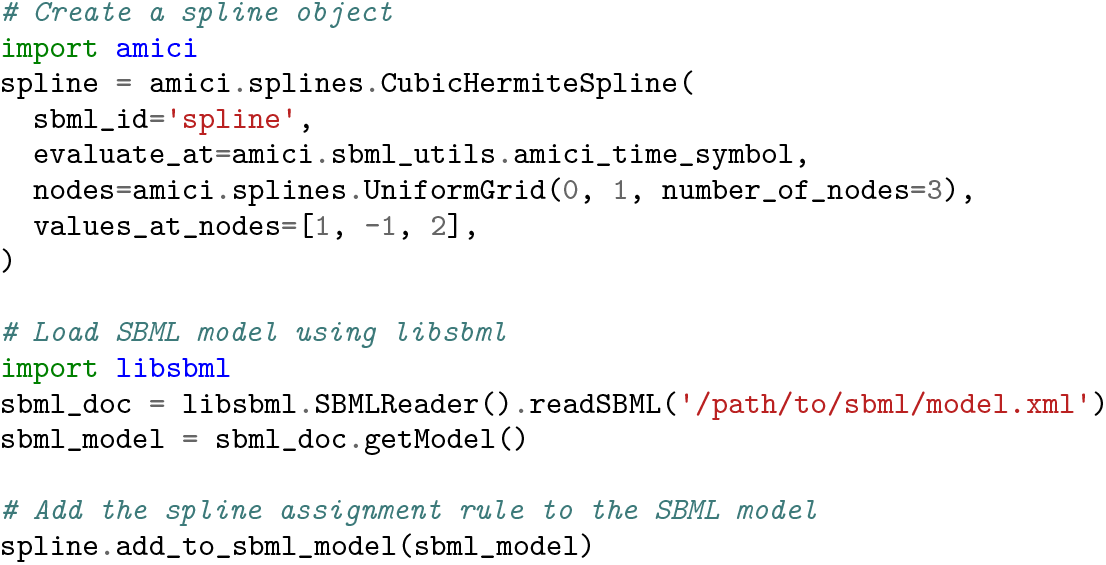
The Python code required to add the spline of Figure 2 to an SBML model.

## 5 Parsing and simulating splines from an SBML model

A model containing an assignment rule of the type presented in Section 4 is a valid SBML model. This means SBML-aware simulation tools can read it, ignore the annotations, parse the piecewise MathML expression and simulate the model. While AMICI is able to handle piecewise formulas correctly too, due to the way symbolic computation is carried out when AMICI processes the SBML file, the resulting code generation times are extremely long, making the current version of AMICI unsuited to run such models. This problem is solved when a compact XML representation of the spline is contained in the annotation to the assignment rule. AMICI can then read the annotation, and generate the C**++** code for the spline from it, without having to parse the MathML expression. This leads to vastly improved import times.

To quantify this speed-up, we consider a cubic Hermite spline with *n* equidistant nodes on the interval [0, 1], known values at the nodes randomly drawn and derivatives computed by finite differences. Then, we study, as *n* changes, the time required by AMICI to read the SBML file and compile it to native code. Results are displayed in Figure 5, showing an exponential dependence of the import time on *n*, independently of whether MathML piecewise expressions or AMICI-specific spline annotations are used. However, the growth rate in the case of piecewise expressions is much larger, making it clear that splines with more than a few dozens of nodes are unfeasible, especially in the case of iterative model development. Model simulations, on the other hand, take similar time and have similar accuracy for both methods. This is expected, since the mathematical formulas required to compute the polynomial segments are always the same, independently of how the spline is parsed from the SBML file. The C**++** code generated from a piecewise representation is longer and contains more repetitions than the code generated from a spline annotation, but execution time is essentially the same in both cases. We suspect this is due to modern compilers being very good at code optimization.

**Fig. 5.**
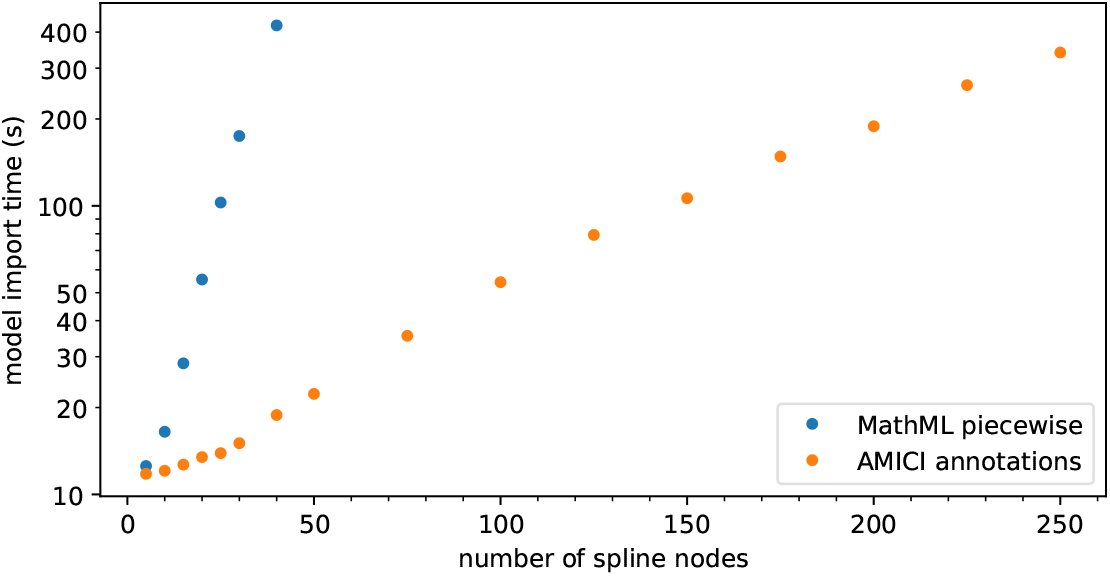
Dependence of model import time on the number of spline nodes. Timing results using MathML piecewise expressions and AMICI annotations are both shown (6 replicates).

## 6 A real-world example

As a real-world example, we consider a chemical reaction network model for the JAK2-STAT5 signaling pathway proposed by [19], in which the dynamics of the system depend on a continuous input function. This input function is only measured (with noise) at a finite number of time points, but its value at all intermediate time points is required to simulate the model. In [17] a 5-node interpolating cubic spline was used to model the logarithm of the input function (the logarithm is used in order to ensure positivity). We closely follow their approach, except that we model the logarithm of the input function using a cubic Hermite spline.

While in [17] the spline had only a few degrees of freedom, in order to show what can be achieved with our implementation we consider two different families of splines with higher expressivity: (i) a spline with 15 nodes where derivatives are estimated by finite differences; (ii) a spline with 5 nodes with derivatives fitted alongside the values at the nodes. In both cases, the derivative at the rightmost node is set to zero since the input function appears to have reached a steady state. In order to reduce overfitting, we follow [17] and add a regularization term consisting of the squared L2 norm of the curvature of the spline, which can be easily computed using the AMICI Python library. Since our splines are more expressive than the one used in [17], the risk of overfitting is going to be larger.

Figure 6 shows the best fits obtained for different values of the regularization strength *λ*. The most appropriate value for *λ* can be chosen by comparing the sum of the squared normalized residuals (which is approximately *χ*^2^-distributed) with its expected value [17]; this results in *λ ≈* 75 for the 15-node spline and *λ ≈* 175 for the 5-node spline.

**Fig. 6.**
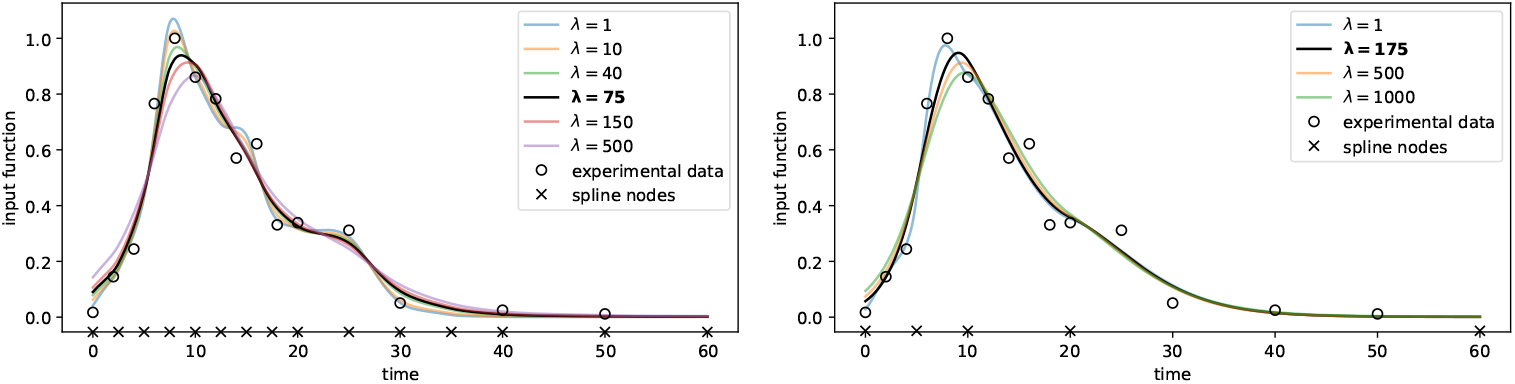
Estimated input function of the JAK2-STAT5 model. The left panel shows the results for a 15-node cubic Hermite spline with derivatives estimated by finite differences, while the right panel uses a 5-node spline with derivatives fitted alongside the other parameters. The squared L2 norm of the input function curvature is used as a regularizer and the estimates for different values of the regularization strength *λ* are plotted. The value of *λ* that results in the sum of the squared normalized residuals being nearest to its expected value is highlighted in bold in the legend.

## 7 Discussion

In conclusion, the new annotation-based spline implementation in AMICI allows to integrate complex spline functions in SBML models without having to spend an inordinate amount of time on code creation. While other tools do not directly benefit from the AMICI-specific annotations, being AMICI one of the few available tools implementing adjoint sensitivity analysis coupled with extensive support of SBML features, we believe this addition will be of use to many members of our community. However, even users of other simulation tools can benefit from using our utility functions to create spline formulas and add them to SBML models, since existing methods only work with numeric data and support fewer spline types. Finally, we believe the SBML standard would greatly benefit from integrating native support for splines and other interpolating functions using a human-readable and easy-to-edit syntax; our XML representation could be a starting point for a discussion among the stakeholders of the SBML ecosystem.

Finally, we remark that, while the main focus of the SBML modelling language is systems biology, it can be used to describe arbitrary systems of differential equations. This allows for the spline implementation described in this paper to be used in other application domains too. For example, AMICI splines have already been used in the epidemiological model proposed by [2] to model the time-dependent effect of non-pharmaceutical interventions on the transmission rate of COVID-19. Thus, we believe this feature will be very useful to mathematical modellers in general, enabling them to increase the expressivity of their models through highly-resolved discretizations of input functions.

## Code availability

These enhancements have been added to AMICI 0.18 [6]. More examples on how to add different kind of splines to SBML models (including the import time benchmark) can be found at https://github.com/AMICI-dev/AMICI/blob/0fc51c0ddcac0661a83931e8b1c4a59f0c6db85c/python/examples/example_splines/ExampleSplines.ipynb, while the code for reproducing the example of Section 6 can be found at https://github.com/AMICI-dev/AMICI/blob/0fc51c0ddcac0661a83931e8b1c4a59f0c6db8python/examples/example_splines_swameye/ExampleSplinesSwameye2003.ipynb.

## 8 Appendix specification for the compact XML spline representation

Our compact XML representation of a cubic spline consists of a `<spline>` element with the following attributes:

- `spline method`, the family of splines to use. At the moment, only `“cubic hermite”` is supported by AMICI, but support for other families (such as monotone splines) may be added in the future.
- `spline bc` (optional, defaults to `“no bc”`), the boundary conditions for the spline, used when derivatives are computed by finite differences. It can be a single value or a tuple in case the boundary condition on the left side differs from the one on the right side. Currently supported values are: `“no bc”` (one sided finite differences are used at the boundary), `“zero derivative”` (first derivative is zero at the boundary), `“natural”` (second derivative is zero at the boundary), `“periodic”` (for a periodic spline).
- `spline extrapolate` (optional, defaults to `“no extrapolation”`), the ex trapolation method for the spline, used to compute the spline value before the first node or after the last one. It can be a single value or a tuple in case the extrapolation method on the left side differs from the one on the right side. Currently supported values are: `“no extrapolation”` (an error should be raised if an attempt is made to evaluate the spline outside its domain), `“constant”` (constant extrapolation, using the value at the boundary), `“linear”` (linear extrapolation, using value and derivative at the boundary), `“polynomial”` (the polynomial segment nearest to the boundary is used outside the domain too), `“periodic”` (for a periodic spline).
- `spline logarithmic parametrization` (optional, defaults to `“false”`), when `“true”` interpolation is carried out in logarithmic scale, in order to enforce positivity of the function (which is no longer a spline in linear scale). Note that spline values and spline derivatives are still specified in linear scale.

The `<spline>` element must also contain the following children elements:

- `<spline evaluation point>`, an element whose single child is the point at which the spline is evaluated, as a MathML expression. In many cases this is simply the model time, as in Figure 2, and at the moment this is the only choice supported by the efficient C**++** implementation in AMICI. However, the tools for creating annotations generate valid MathML expressions also for other choices of evaluation point, so that they can be used with other tools.
- `<spline grid>`, an element whose children are the spline nodes, as MathML expressions.
- `<spline uniform grid>` (if present, replaces `<spline grid>`), a compact representation of a uniform grid of nodes. Its children are the first node, the last node and the step size, all as MathML expressions.
- `<spline values>`, an element whose children are the values of the spline at the nodes, as MathML expressions.
- `<spline derivatives>` (optional, if missing it is computed using finite differences and boundary conditions), an element whose children are the values of the first derivative of the spline at the nodes, as MathML expressions.

